# Interpreting Null Models of Resting-State Functional MRI

**DOI:** 10.1101/2021.03.30.437514

**Authors:** Raphaël Liégeois, B. T. Thomas Yeo, Dimitri Van De Ville

## Abstract

Null models are necessary for assessing whether a dataset exhibits non-trivial statistical properties. These models have recently gained interest in the neuroimaging community as means to explore dynamic properties of functional Magnetic Resonance Imaging (fMRI) time series. Interpretation of null-model testing in this context may not be straightforward because (i) null hypotheses associated to different null models are sometimes unclear and (ii) fMRI metrics might be ‘trivial’, i.e. preserved under the null hypothesis, and still be useful in neuroimaging applications. In this commentary, we review several commonly used null models of fMRI time series and discuss the interpretation of the corresponding tests. We argue that, while null-model testing allows for a better characterization of the statistical properties of fMRI time series and associated metrics, it should not be considered as a mandatory validation step to assess their relevance in neuroimaging applications.

## Introduction

Functional interactions in the resting human brain are organized in complex spatio-temporal patterns (Greicius et al., 2003; Smith et al., 2013). The classical way of characterizing these patterns is within *static* modelling frameworks, e.g., evaluating functional connectivity (FC) from the correlation between whole-run functional Magnetic Resonance Imaging (fMRI) time series across different brain regions (Biswal et al., 1995; Zalesky et al., 2010; Margulies et al., 2016). In contrast, recent work has brought out promises of considering *dynamic* frameworks, such as sliding window methods (Sakoğlu et al., 2010; Handwerker et al., 2012; Allen et al., 2014), to account for temporal variability of connectivity patterns (see Preti et al. (2017); Lurie et al. (2020) for recent reviews). Such dynamic FC frameworks raise questions like *‘Are transition probability patterns between brain states different in the original data and in surrogate data with matched spatial covariance struc-12 ture?’ or ‘Are original fluctuations of a metric only due to autocorrelation of fMRI time series or are they revealing the presence of more complex (e.g., nonlinear) functional interactions?’*. These questions can be addressed using null-model testing that consists in comparing the original data vs. random data preserving a subset of original statistical properties, a.k.a. surrogate data (e.g., Theiler et al., 1992).

Null-model testing has been widely used in dynamic FC studies with apparently contradictory 18 outcomes: while most studies concluded the rejection of the null hypothesis (Chang and Glover, 2010; Handwerker et al., 2012; Zalesky et al., 2014), others reported difficulties in rejecting the null hypothesis (Hindriks et al., 2016; Laumann et al., 2016; Liégeois et al., 2017). This observation is the starting point of our commentary. We first recall basics of null-model testing and provide a detailed description of popular null models of fMRI time series such as phase randomization and the autoregressive null model. As we will see, these null-model methods are related but test for distinct null hypotheses that need to be carefully understood in order to interpret testing outcomes. Then, we argue that a statistical metric preserved under a null hypothesis can still be useful in neuroimaging applications, thereby making null-model testing of limited relevance in this context. Finally, we discuss a few concrete examples and provide recommendations on the use and interpretation of null-model testing in dynamic FC studies.

### Basics of null-model testing

We consider a time-series dataset **D** with dimensions *N* × *T* where *N* is the number of variables and *T* is the number of time points, and we denote **d**_*t*_ as the *N* × 1 vector encoding values of the *N* variables at time *t*. Without loss of generality, we further assume that each of the *N* time courses composing **D** are centered and of unit variance. The autocorrelation sequence of **D** is defined as the following sequence of *N* × *N* matrices:

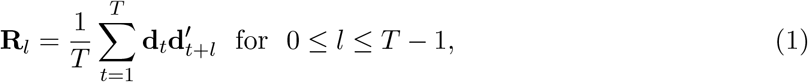

where ′ denotes transpose. Diagonal elements of **R**_*l*_ encode autocorrelation within individual time courses, while off-diagonal terms of **R**_*l*_ encode autocorrelation between pairs of time courses. Note that **R**_0_ encodes the classical correlation of **D** (or, equivalently, its covariance since time courses are assumed to be of unit variance) and we refer to it as Σ hereafter.

The goal of null-model testing is to gain insights into the statistical nature of **D** by testing the validity of a null hypothesis 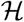 to represent **D**. Null-model testing frameworks typically consist of the three following steps.

1. Generate data preserving a subset of original statistical properties of **D**, but that are otherwise random. This data is referred to as *surrogate* data and we denote it **D**_*s*_. Importantly, **D**_*s*_ comprises several data instances and the method used to generate **D**_*s*_ defines the null hypothesis 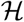.
2. Compute a metric *m* in both the original and surrogate datasets. The metric value in the original dataset is denoted as *m*_*o*_, and the metric values computed from each surrogate data instance in **D**_*s*_ define a null distribution denoted 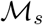.
3. Compare *m*_*o*_ and the null distribution 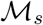. If *m*_*o*_ is different from 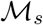, the null hypothesis 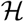 is not sufficient to fully describe **D**. If *m*_*o*_ is not different from 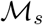, the null hypothesis 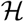 cannot be rejected.

#### A toy example

Consider the case of a univariate dataset **D** with 50 time points, i.e., *N* = 1 and *T* = 50, shown in Figure 1A. To better characterize **D**, let us apply two null-model approaches.

**Figure 1:**
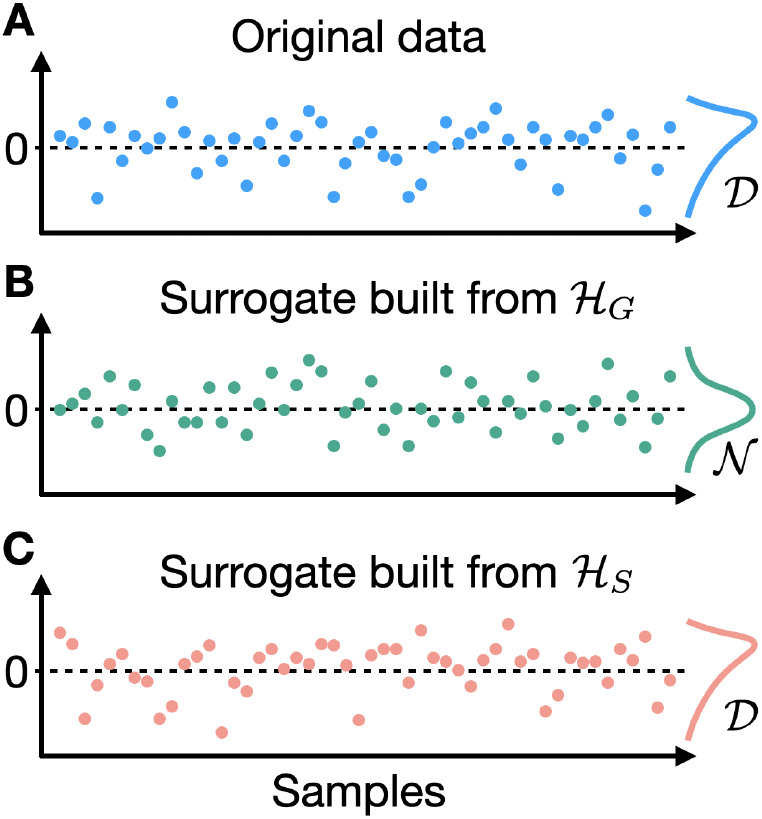
Comparing original and surrogate data. (A) Original data containing 50 time points with sample distribution 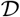. (B) One instance of surrogate data generated using a Gaussian univariate null hypothesis 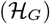. The corresponding (Gaussian) sample distribution is denoted 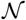 with mean and standard deviation matched to the origi-nal data. (C) One instance of surrogate data generated by shuffling original data. The null hypothesis corresponding to this procedure is 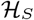 and the corresponding sample distribution is the same as the original data 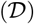.

First, we consider the null hypothesis, denoted 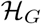, that **D** can be represented by independent and identically distributed (i.i.d.) Gaussian random variables. Step 1 in the above procedure consists in producing surrogate data **D**_*s*_ from an i.i.d. Gaussian distribution with length, mean (*µ*) and standard deviation (*σ*) matched to the length, mean and standard deviation of **D** (Theile et al., 1992). An instance obtained following this procedure is shown in Figure 1B. Then, in Step 2 one needs to compute a metric *m* in the original and surrogate data that is not directly defined by *μ* and *σ* as these quantities are by construction equal in **D**_*s*_ and **D**. One could for example compare the number of time points above a given threshold in both the original and all surrogate datasets, leading to *m*_*o*_ and 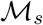. Finally, in Step 3 values of *m*_*o*_ and 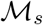 are compared in order to evaluate validity of the null hypothesis. In this case the null hypothesis, denoted 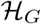, is straightforward and could be expressed as:

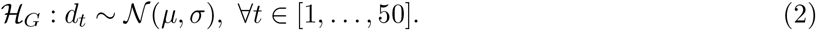

We will see in the next null-model framework that the null hypothesis defined by Step 1 is not always easy to identify. Then, the interpretation of the comparison performed in Step 3 depends on the null hypothesis: finding that *m*_*o*_ is different from 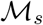 suggests that **D** contains statistical information beyond *μ* and *σ*, whereas finding that *m*_*o*_ is not different from 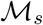 indicates that 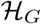 cannot be rejected.

The second null-model framework we consider consists in generating surrogate data of Step 1 by shuffling time points of **D**, as illustrated in Figure 1C (Scheinkman and LeBaron, 1989). The corresponding null hypothesis, denoted 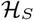, is not explicitly defined by this procedure and could be expressed as:

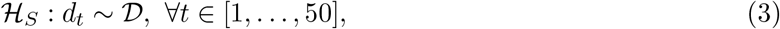

where 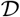 is the sample distribution of **D** (Figure 1A). As such, **D**_*s*_ preserves the mean and standard deviation of **D** but also all other statistics defined by 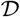 such as kurtosis and skewness. The metric computed at Step 2 needs to be chosen accordingly: for example, comparing the number of time points above a given threshold in **D** and **D**_*s*_ would in this case be meaningless since this quantity is by construction equal in the original and surrogate datasets. An appropriate metric *m* could be, e.g., the average autocorrelation in **D** and **D**_*s*_. Finally, comparison of *m*_*o*_ and 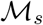 in Step 3 provides information on validity of 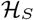: finding that *m_*o*_* is different from 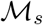 shows there is autocorrelation in **D** whereas finding that *m*_*o*_ is not different from 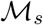 suggests there is no temporal ordering information in **D**.

Beyond illustrating main steps of null-model testing frameworks, several important observations can be made from this simple toy example. First, the null hypothesis associated to the surrogate data generation procedure (Step 1) is not always straightforward to identify. Second, it is crucial to clearly identify this null hypothesis in order to (i) define a relevant metric *m* to be compared in original and surrogate datasets (Step 2) and (ii) properly interpret the result of this comparison (Step 3). Third, null hypothesis rejection does not have a unique interpretation as it can be due to violation of any property preserved by the null. For example, rejecting 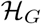 could be due to non-Gaussianity, non-stationarity, the presence of autocorrelation in **D**, or a combination of these. A careful choice of the metric *m* in Step 2 might in certain cases allow to narrow down causes of null-model rejection, i.e., in this case determining which statistical property among non-Gaussianity, non-stationarity, or autocorrelation, led to the rejection of 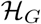. The description of the different metrics types is beyond the scope of this paper and we refer the reader to the excellent review by Lancaster et al. (2018) for a detailed presentation of these metrics and the statistical properties they capture. Finally, note that not rejecting the null hypothesis at Step 3 using a given metric *m*does not imply that the null model is sufficient to fully describe **D**.

### Null models of fMRI time series

The two null models presented in the above toy example are not suited to explore the statistical properties of fMRI time series because the latter are known to be autocorrelated, leading to an -uninformative- rejection of 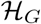 and 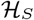. We present two null-model frameworks that account for this autocorrelation while also preserving fMRI covariance structure: multivariate autoregressive null models and phase randomization (Theiler et al., 1992).

Multivariate autoregressive (AR) null models assume that **D** can be represented as an autoregressive model. We focus on the case of first-order AR (AR-1) models, i.e., the null hypothesis, denoted 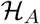, is that data at time *t* can be expressed as a weighted sum of data at time *t* − 1 plus Gaussian noise:

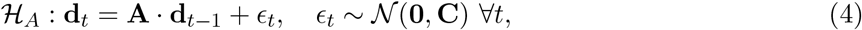

where **A** is a model parameter of size *N* × *N* and *ϵ*_*t*_ is i.i.d. Gaussian noise. Each null sample in **D**_*s*_ is initialized by randomly selecting a time point in **D**, and then generating *T* − 1 new time points by successively applying Eq. (4), with **A** and **C** identified from the original data. It can be shown from the Yule-Walker equations that **D** and **D**_*s*_ have the same correlation (Σ) and first-order autocorrelation (**R**_1_, see Eq. (1)) structures (Yule, 1927; Walker, 1931). More details on statistical properties of AR null models and their application to fMRI time series are found in Liégeois et al. (2017).

Phase randomization (PR) is another method that has been widely used to explore properties of fMRI time series (e.g., Handwerker et al., 2012; Hindriks et al., 2016). In this framework, surrogate data of Step 1 is generated by performing Discrete Fourier Transform (DFT) of each time course, adding a uniformly distributed random phase to each frequency, and then performing the inverse DFT (Prichard and Theiler, 1994). Importantly, the random phases need to preserve Hermitian symmetry and are generated independently for each frequency, but they are the same across the *N* variables, i.e., brain regions or voxels in this case. From the Wiener-Khintchine theorem, it follows that surrogate data generated by PR preserves the full autocorrelation structure of **D** defined in Eq. (1) (Wiener, 1930; Khintchine, 1934). As such, the corresponding null hypothesis is that **D** can be represented using a linear and Gaussian model:

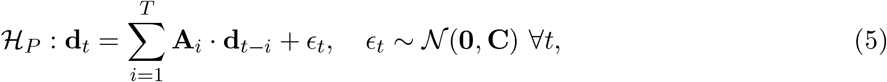

where **A**_*i*_ are model parameters of size *N* × *N* and *ϵ*_*t*_ is i.i.d. Gaussian noise. It can be seen by comparing Eqs. (4) and (5) that while both the AR-1 null and PR assume a linear and Gaussian representation of **D**, PR preserves more statistical information from the original data. Indeed, the AR-1 null preserves Σ and **R**_1_ whereas PR preserves Σ and the whole autocorrelation sequence **R**_1_,…, **R**_*T*_ defined in Eq. (1). We also note that while the model parameter **A** can be identified in Eq. (4) provided at least *N* + 1 time points are available in **D** (i.e., *T* > *N*), PR model parameters **A**_*i*_ for *i* = 1,…, *T* in Eq. (5) cannot be explicitly identified from the autocorrelation sequence of **D** (Stoica and Moses, 2005). Despite these practical differences, both approaches were found to yield similar conclusions about the nature of fMRI time series which suggests that **R**_1_ captures a significant proportion of fMRI autocorrelation structure (Liégeois et al., 2017).

The main properties of the four null models introduced in this paper are summarized in Table 1. The first two frameworks corresponding to 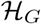 and 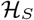 do not preserve any temporal structure of time series data, thereby leading to rejection of the corresponding null hypotheses when applied to fMRI time series. We have included them in order to provide a complete description of the links between popular null-model frameworks, while also motivating the use of PR and the multivariate AR null in this context.

**Table 1:** Main properties of the four null-model frameworks presented in this commentary. For each framework, the corresponding null hypotheses and statistical properties of **D** being preserved in **D**_*s*_ are reported. A non-exhaustive list of valid measures *m*, i.e., measures that are not entirely defined by the statistical properties shared by *D* and *D*_*s*_, to be computed in Step 2 are also provided. 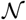 i.i.d. Gaussian distribution - 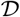: sample distribution of *D* - *κ*: kurtosis - Σ: correlation matrix - **R**_*p*_: *p*^*th*^ order autocorrelation matrix as defined in Eq. (1).

|  | Gaussian<br>and i.i.d. | Data<br>shuffling | AR-1<br>Randomization | Phase<br>Randomization |
| --- | --- | --- | --- | --- |
| Null<br>hypothesis | $\mathcal{H}_G$:<br>$\mathbf{d}_t = \epsilon_t, \epsilon_t \sim \mathcal{N}$ | $\mathcal{H}_S$:<br>$\mathbf{d}_t = \epsilon_t, \epsilon_t \sim \mathcal{D}$ | $\mathcal{H}_A$:<br>$\mathbf{d}_t = \mathbf{A} \cdot \mathbf{d}_{t-1} + \epsilon_t$ | $\mathcal{H}_P$:<br>$\mathbf{d}_t = \sum_{i=1}^T \mathbf{A}_i \cdot \mathbf{d}_{t-i} + \epsilon_t$ |
| Properties<br>preserved | $\Sigma$ | $f(\mathcal{D}) : \Sigma, \kappa, \text{etc.}$ | $\Sigma, \mathbf{R}_1$ | $\Sigma, \mathbf{R}_1, \dots, \mathbf{R}_T$ |
| Examples<br>of useful $m$ | • Kurtosis ($\kappa$)<br>• Nb. of points<br>> threshold<br>• $\mathbf{R}_1$ | • $\mathbf{R}_1$<br>• Measures of<br>nonlinearity or<br>nonstationarity | • $\mathbf{R}_p$ with $p > 1$<br>• Kurtosis ($\kappa$)<br>• Measures of<br>nonlinearity | • Measures of<br>nonlinearity<br>• State transition<br>patterns |

#### Related models and methods

PR and the AR null are most commonly used to explore properties of measures of interest computed from fMRI time series, but other methods have been considered. For example, Chang and Glover (2010) used a variation of the AR null that, instead of identifying a single multivariate AR model for the whole *N*-variate system of fMRI time series as described in Eq. (4), generates AR surrogates for each pair of variables separately. As a result, bivariate surrogate time series are independent for each pair of variables and do not account for potential higher-order spatial interactions present in fMRI time series, thereby causing rejection of the corresponding null hypothesis (e.g., Figure 6 in Liégeois et al., 2017). Then, a variant of PR in which the random phase added to the DFT components of a given frequency is different for each variable has been introduced recently (Abrol et al., 2017). This framework, referred to as ‘inconsistent’ PR, does not preserve the correlation structure of **D** which causes rejection of the corresponding null hypothesis when applied to fMRI time series. Another recent framework produces surrogate data that preserves original correlation structure as well as the power spectral density of each individual time course, which by the Wiener-Khintchine theorem amounts to preserving the autocorrelation structure within each time course (Laumann et al., 2016). In other words, this framework preserves original Σ as well as the diagonal entries of the autocorrelation sequence **R**_1_,…, **R**_*T*_ of **D** and could therefore be considered as an intermediate between the Gaussian i.i.d. null and PR in terms of statistical properties being preserved. Finally, other approaches factor in underlying anatomical information (Pirondini et al., 2016; Petrovic et al., 2020) or account for more complex statistical properties (e.g., Theiler et al., 1992; Breakspear et al., 2003; Van De Ville et al., 2004).

### Statistically ‘trivial’ does not mean useless

When evaluating the relevance of a dynamic fMRI (or FC) metric, it has become common practice to require that this metric produces different results in original and surrogate data in order to ensure it does not reflect ‘trivial’ statistical data properties (see, e.g., Preti et al., 2017, and references therein). The word ‘trivial’ here refers to statistical properties preserved by the null-model framework applied to fMRI time series, usually multivariate AR or PR. While we agree these statistical properties are to some extent ‘trivial’ as 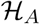 or 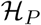 only preserve linear and Gaussian fMRI features, we argue that rejecting (or not rejecting) these null hypotheses is uninformative about the neurological or clinical use of the considered fMRI metric.

To this end, let us reformulate the interpretation of Step 3 testing outcome using the representation of Figure 2. The outer green circle represents the set of all statistical properties characterizing original data **D**, i.e., all spatial, temporal, and spatiotemporal statistical moments of **D**. The three inner circles represent statistical information preserved by three different null frameworks: Gaussian and i.i.d., AR-1, and PR. We did not include data shuffling for simplicity purposes as the corresponding null hypothesis is not nested within the other hypotheses, i.e., 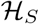 contains strictly more information than 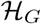 but is not contained in 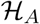.

**Figure 2:**
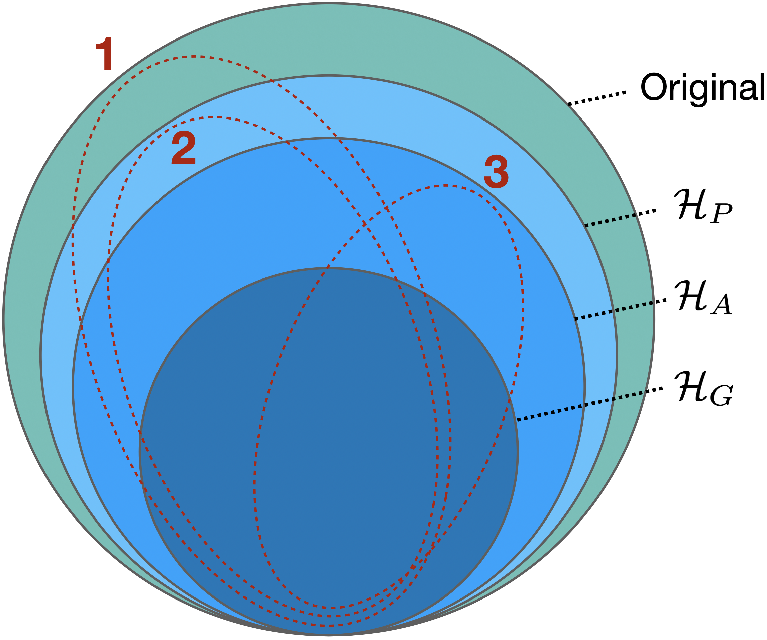
Nested structure of original statistical properties (outer green circle) being preserved by 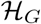 (Gaussian i.i.d. null model, inner dark blue circle), 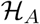 (AR-1 null, intermediate medium blue circle), and 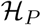(phase randomization, intermediate light blue circle). Red dotted ellipses represent different subsets of statistical information used in the practical examples at the end of the paper.

Rejecting a null hypothesis at Step 3 shows that the metric being considered exploits statistical properties *beyond* the ones preserved by the null hypothesis. For example, observing that an fMRI metric is different in original and AR-1 surrogates suggests that this metric exploits statistical information beyond 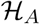 i.e., beyond fMRI correlation (Σ) and first-order autocorrelation (**R**_1_) structures (e.g., this could be the case of a metric exploiting statistical information of curves 1 and 2 in Figure 2). Framed differently, the metric exploits some forms of nonlinearity, non-Gaussianity, non-stationarity, or higher-order autocorrelation of fMRI time courses. Importantly, exploiting fMRI statistical information beyond Σ and **R**_1_ is not informative about relevance of that metric in neuroimaging applications such as disease classification, fingerprinting, etc. Indeed, there is no reason for a metric outside of 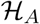, e.g., a measure of nonlinearity such as kurtosis, to be *a priori* more relevant than a metric that lies within 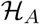 -and therefore called ‘trivial’-such as average first-order autocorrelation. In sum, the statistical information captured by fMRI or FC metrics and assessed using the null-model frameworks presented in this paper is unrelated to their clinical use which should be tested within a separate framework.

As an illustration, consider the case of fMRI time series correlation (Σ) that is classically used to evaluate FC. Utility of this metric in a wide range of neuroimaging applications no longer needs to be proven (e.g., Buckner et al., 2013, and references therein), and any metric derived from Σ such as partial correlations (Ryali et al., 2012) or graph metrics computed from Σ (Meunier et al., 2010; Mišićc et al., 2016) and Σ^−1^ (Liégeois et al., 2020) would be found to be the same in original and surrogate data generated from 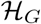, 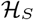, 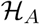 or 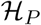 as all these null hypotheses preserve the original Σ (Table 1). This further illustrates that ‘trivial’ statistical information preserved within a null-model framework can be useful in neuroimaging applications. Note that metrics that are statistically equivalent and hence preserved under the same null-hypotheses, such as Σ^−1^ and Σ, can be of different usefulness; e.g., FC evaluated from *partial* correlations (derived from Σ^−1^) was shown to outperform FC evaluated from correlations in several prediction tasks (Dadi et al., 2019). A similar rationale applies for measures derived from the autocorrelation sequence of fMRI time series that is preserved by 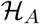 and 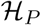, and was shown to provide insight into brain functional organization (Mitra et al., 2014) and its link with human behaviour (Liégeois et al., 2019).

#### A measure of model complexity

In the above we argue that null-model testing outcomes are not informative about relevance of a given metric in neuroimaging applications. This being said, these tests provide insight into the statistical complexity of the metric which could be useful in terms of model selection. For example, finding that a metric is shown to be equivalent in original and AR-1 surrogates suggests that it is exploiting statistical properties preserved by 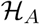: Σ and **R**_1_. These two parameters can be represented by *N* · (3*N* − 1)/2 parameters (*N*^2^ for **R**_1_ and *N* · (*N* − 1)/2 for Σ) which gives an upper bound on the parametric complexity of the metric. As for PR, the full autocorrelation sequence preserved by 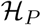 is described by ≈ *T* · *N*^2^/2 parameters^1^. This highlights that the notion of ‘trivial’ is relative: while PR only preserves linear and Gaussian properties, the parametric space defined by such models can get considerable when considering typical fMRI time series with *N* ≈ 100 and *T* ≈ 1000. We show in the first practical example hereunder how these consideration can be included to perform model selection of fMRI time series.

### A null hypothesis can (almost) always be rejected

Rejecting a null hypothesis means that the corresponding null model is not sufficient to fully describe **D**. This is correct, but should not be over-interpreted. As an illustration, consider the distribution of US women height obtained from the US Census Bureau^2^ and represented by the grey bars in Figure 3.

**Figure 3:**
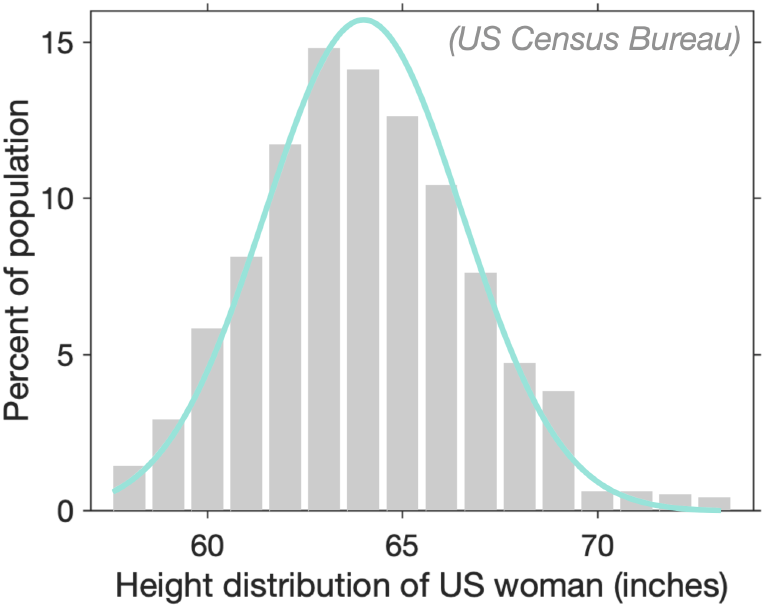
Data models are imperfect. Grey bars represent original distribution of height of US women aged 30-39 and the green line represents a Gaussian distribution matched to original data. A Gaussian model does not account for tall women (> 80 *in*), yet it provides a useful representation of US women height distribution.

Let us evaluate the relevance of a Gaussian representation of this distribution using null-model testing. Step 1 consists in identifying a Gaussian model from the data and generating surrogate data from it. The corresponding null hypothesis 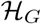 is characterized by a mean *μ* (in this case *μ* = 64 *in*) and a standard deviation *σ* (in this case *σ* = 2.6 *in*). Let us now define *m* used in Step 2 as the number of women taller than 80 inches. Under the Gaussian null, probability of finding a woman meeting this criterion is *p* = 3.8 · 10^−10^ which leads to rejection of the Gaussian null at Step 3 since *m*_*o*_ is larger than the null distribution 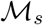(i.e., the number of women taller than 80 *in* is in reality non-zero). In other words, the Gaussian null is rejected because it does not fully capture properties of the data, in this case the proportion of extreme values of US women heights. Yet, it seems reasonable to consider that a Gaussian model of the data as represented by the green line in Figure 3 is informative about US women height distribution.

Two conclusions shall be drawn from this example. First, it seems virtually always possible to choose a metric *m* at Step 2 that leads to rejection of the null hypothesis by focusing on known peculiarities of the original dataset, e.g., the distribution of extreme values as illustrated above. Second, null hypothesis rejection does not imply that the corresponding model of the data is useless. Interpretation of this result should be performed in light of the more general model selection question, as illustrated in the first practical example hereunder.

### Practical examples

We present three practical examples inspired from the dynamic FC literature. While we aimed to consider realistic applications and results, we did not actually perform the analyses as our goal is to discuss the interpretation of results typically encountered in null-model testing analyses of functional dynamics.

#### Example 1: Brain states dynamics

Let us consider a classical sliding window correlation (SWC) approach of fMRI time series followed by a clustering of FC connectivity matrices into so-called brain states. We use a clustering into five states and focus on the transition probability matrix that encodes transition probability between each pair of brain states (e.g., Allen et al., 2014). This FC dynamics summary metric, as well as related measures such as dwell time, were for example used to describe differences between control and schizophrenia groups (Damaraju et al., 2014; Du et al., 2016). Assume we compare the original transition probability matrix of one subject and the distribution of transition probability matrices computed in AR-1 surrogate data, as illustrated in Figure 4A, and find that three matrix entries are different in original and surrogate matrices. This result suggests that the transition probability matrix captures, at least partly, statistical information above and beyond the one preserved by the AR-1 null (e.g., curves 1 or 2 in Figure 2). More precisely, results of Figure 4A suggest that three entries of the transition probability matrix rely upon original statistical information beyond 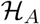, while the 22 remaining entries out of 25 exploit information within 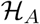.

**Figure 4:**
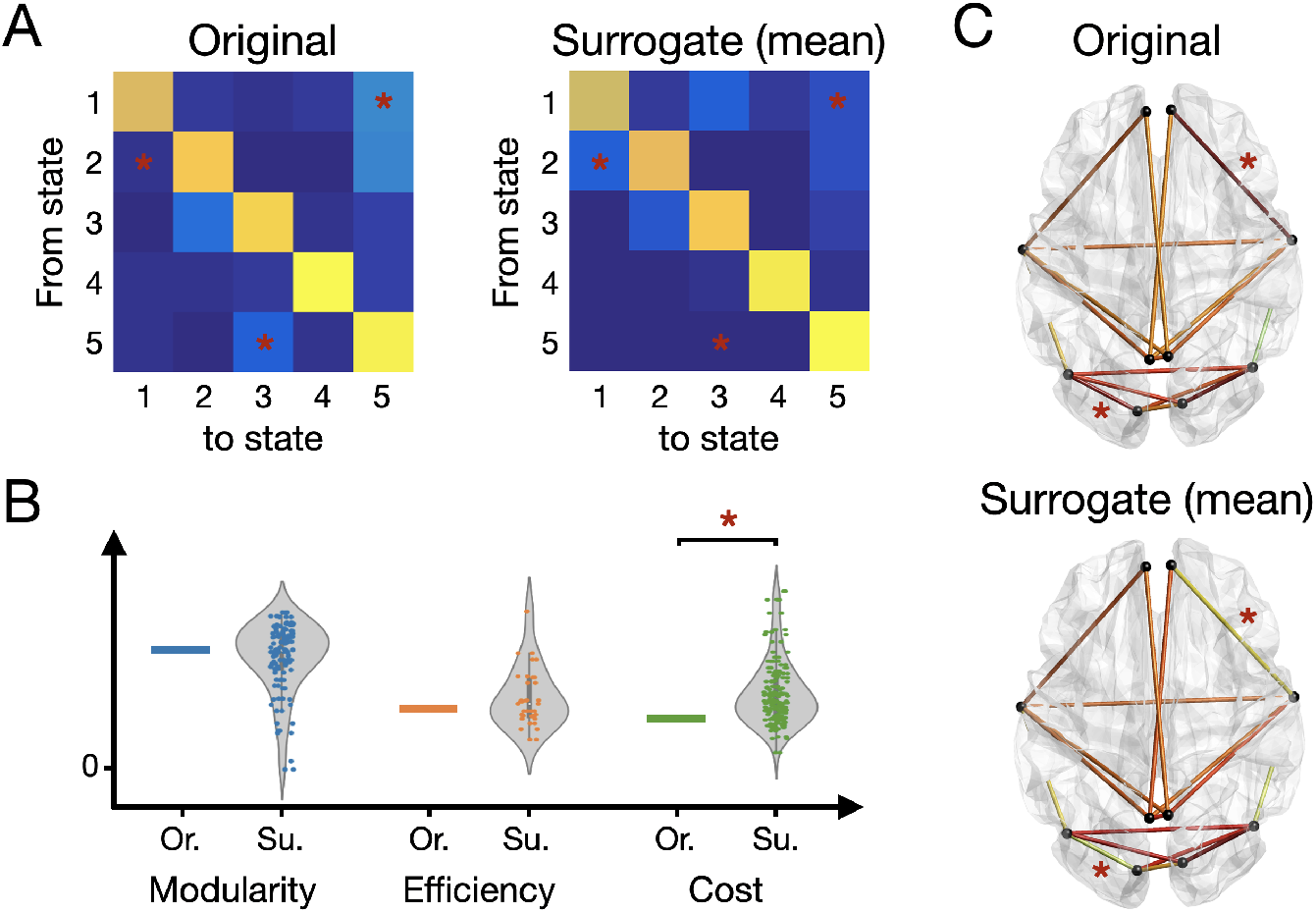
Illustrations of the three practical examples. Red stars denote significant differences between original and surrogate measures. (A) Transition probability matrices between five brain states computed in original (left) and surrogate (right) data. Warmer colors denote higher transition probabilities. Three matrix entries are found to be different between original and surrogate data: transitions probability from state 1 to 5, from state 2 to 1, and from state 5 to 3. (B) Three graph-theoretic measures are computed in original (Or.) and surrogate (Su.) data. All surrogate values are represented (i.e., not only the mean), and the corresponding violin plot is represented. One metric (cost) is found to be higher in surrogate than in original data. (C) Amplitude of sliding window correlation (SWC) fluctuations in a selection of 18 edges pertaining to the visual and default mode networks, in original (top) and surrogate (bottom) data. Two pairs of nodes are found to exhibit different SWC fluctuations in original and surrogate data.

Based on the considerations presented in the above sections, we make the following comments. First, similar to the case discussed in Figure 3, rejecting 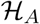 does not disqualify the AR-1 representation of the data. This question has to be considered in a more general model selection framework balancing model fit (in this case, 22 out of 25 entries are reproduced by the AR-1 model) and model complexity. Then, while results show that transition probabilities from state 1 to 5, from state 2 to 1, and from state 5 to 3 reflect more complex (i.e., beyond 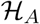) functional dynamics, this is uninformative about neurological relevance or clinical use of these matrix entries as compared to other matrix entries. Finally, considering five brain states corresponds to using 5*N* (*N* − 1)/2 parameters to summarize functional dynamics whereas the AR-1 representation uses *N* (3*N* − 1)/2 parameters. This increased model complexity could be taken into account when discussing costs and benefits of brain state representations.

#### Example 2: Graph metrics

Let us focus on graph metrics computed from brain state transition networks. These networks are computed by first applying a SWC approach, and then defining a network based on the transitions between successive FC states, as for example proposed in Ramirez-Mahaluf et al. (2020). Three graph metrics are considered: modularity, efficiency, and cost. These metrics are computed in both the original (Or.) and PR surrogate (Su.) data samples, as illustrated in Figure 4B. We assume no significant difference is found between original and surrogate data for modularity and efficiency, while the cost is found to be significantly higher in surrogate data as compared to original data. This suggests that modularity and efficiency reflect linear and Gaussian features of the original time series (e.g., curves 2 or 3 in Figure 2), whereas cost exploits at least partly nonlinear or non-Gaussian features (e.g., curve 1 in Figure 2). As in the previous example, this result is uninformative about the neuroscientific relevance or clinical use of the three metrics. Finally, note that it would not be relevant to perform a null-model analysis on metrics derived from a functional graph defined by Σ (e.g., Hallquist and Hillary, 2019), since Σ is by construction preserved by both 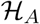 and 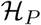.

#### Example 3: Amplitude of SWC fluctuations

Amplitude of SWC fluctuations, formally evaluated from the variance of SWC time series, has also been used to describe functional dynamics (e.g., Liégeois et al., 2016; Hindriks et al., 2016). In this example we focus on the amplitude of SWC in a selection of 18 pairs of brain nodes (i.e., edges) located in the visual and default mode networks, computed both in original and AR-1 surrogate data (Figure 4C). Each edge reflects variance of SWC time series for the two corresponding nodes, with warmer colors denoting higher variance. Results presented in Figure 4C show that among the 18 edges considered in this analysis, no statistical difference between original and surrogate data is found for 16 of them, whereas 2 are found to be different. This suggests that fluctuations of SWC between the nodes corresponding to these two edges reflect, at least partly, nonlinear, non-Gaussian, or higher order linear functional interactions (e.g., curves 1 or 2 in Figure 2). In contrast, functional interactions represented by the 16 remaining edges is a priori capturing statistical information within 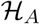 (e.g., curve 3 in Figure 2). Again, this result is uninformative about the neuroscientific relevance or clinical use of functional interactions represented by these two edges, as compared to the remaining 16 edges.

## Conclusions and Recommendations

Null-model testing is widely used in dynamic FC studies. In this commentary we show that the choice of an appropriate null model, as well as the interpretation of the testing outcomes, requires caution. Concretely, we make the following suggestions on the use and interpretation of null-model testing of brain functional dynamics.

– We recommend using the multivariate autoregressive null or phase randomization to generate surrogate data of fMRI time series. It is important to be aware of the corresponding null hypotheses reported in Table 1 in order to interpret testing outcomes.
– Comparing a metric in original and surrogate data is uninformative about clinical or neurological utility of that metric. Therefore, null-model testing should *not* be considered as a mandatory step to validate a novel (dynamic) FC metric.
– Comparing a metric in original and surrogate data provides a better characterization of statistical information exploited by that metric. This can be used, e.g., to assess its parametric complexity.

While highlighting important limitations of null-model testing to explore functional dynamics, these considerations also draw promising avenues for defining new statistical markers of fMRI time series. Indeed, we have seen that statistical measures preserved by AR-1 or PR null models are a priori as relevant as more complex metrics to describe functional dynamics. In this context, fMRI time series’ correlation has received a lot of attention as a marker of functional connectivity, but quite surprising is the fact that ‘next circles’ of statistical properties preserved by linear and Gaussian null models represented in Figure 2 i.e., their autocorrelation structure, has received limited attention so far.

Overall, this commentary provides the theoretical background to design and interpret null-model testing of brain functional time series. Beyond warning about potential misuse in dynamic FC studies, we highlight promises of this framework to identify novel metrics exploiting simple and interpretable features of functional dynamics.

## Acknowledgements

RL acknowledges support by the Swiss National Centre of Competence in Research - Evolving Language (grant number 51NF40 180888).

## Footnotes

1 The 1/2 factor comes from the fact that PR assumes periodic time series (Theiler et al., 1992).Therefore, 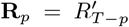 for 0 ≤ *p* ≤ *T*, and only half of the autocorrelation sequence can be described by independent parameters. The exact number of independent parameters depends on *T* being odd or even.

2 https://www2.census.gov/library/publications/2010/compendia/statab/130ed/tables/11s0205.pdf

